# Sensitive detection of circular DNAs at single-nucleotide resolution using guided realignment of partially aligned reads

**DOI:** 10.1101/654194

**Authors:** Iñigo Prada-Luengo, Anders Krogh, Lasse Maretty, Birgitte Regenberg

## Abstract

Circular DNA has recently been identified across different species including human normal and cancerous tissue, but short-read mappers are unable to align many of the reads crossing circle junctions and hence limits their detection from short-read sequencing data. Here, we propose a new method, Circle-Map, that guides the realignment of partially aligned reads using information from discordantly mapped reads. We demonstrate how this approach dramatically increases sensitivity for detection of circular DNA on both simulated and real data while retaining high precision.

## Main

Circular DNA from all parts of the genome has recently been discovered in model organisms ^1,2^, as well as both normal ^3,4^ and cancerous human tissues ^5,6^ using short-read sequencing based approaches. The primary circular DNA signals in short read data are “discordantly” mapped paired-end reads and “split-reads” crossing the circle breakpoint. Yet standard short read aligners do not reliably detect the latter signal, which is vital for determining the exact circle coordinates, as only alignments that are collinear with the reference sequence are typically considered. Reads that cross the breakpoint of a circular DNA will therefore typically be reported either as two or more separate alignments (a primary and some supplementary alignments) when the read has long “anchors” on both sides of the breakpoint or with a major aligned part and a minor, “soft-clipped” (i.e. unaligned) part.

We have developed a new bioinformatics method to accurately identify circular DNA breakpoints called Circle-Map. The overall idea in this method is to use information from discordantly mapped paired-end reads as a *prior* for realigning the soft clipped parts of breakpoint reads, which in turn should allow for more accurate detection of circle breakpoints. Circle-Map consists of three main steps: 1) circular DNA candidate read identification, 2) breakpoint graph construction and 3) soft-clip realignment (Fig. 1).

**Figure 1.**
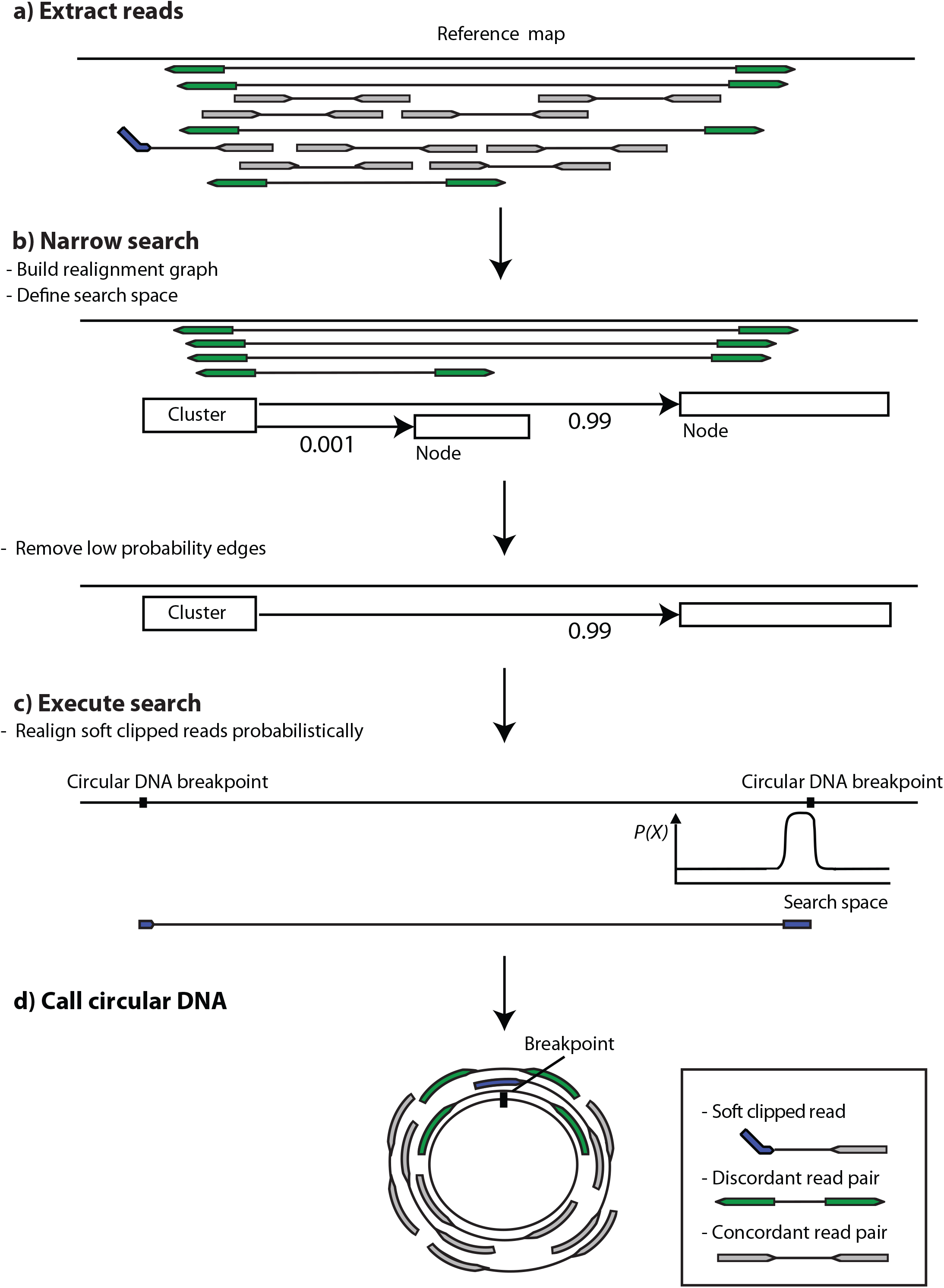
Circle-Map read realignment strategy. **(a)** Reads are mapped to the reference genome and discordantly aligned reads (green) and alignments containing soft clips (blue) are extracted; concordantly aligned reads (grey) are ignored. **(b)** Using the extracted reads, a graph of putative breakpoint connections between genomic regions is constructed and used as a prior to narrow down the genomic search space for realigning soft clipped reads. **(c)** Non-aligned parts of the soft clipped reads are realigned probabilistically using the breakpoint graph as guide. **(d)** Evidence from split-reads and discordant reads are combined to create the final circle calls together with information about concordant, split-read and discordant read coverage for each circle.

### Candidate read identification

Circle-Map workflow begins by performing an initial pass through a query name sorted alignment file to extract reads potentially originating from circular DNA (“circular DNA candidate reads”): discordant read pairs, soft-clipped reads and hard clipped reads (Fig. 1a). For every read pair, Circle-Map labels the pair as discordant if the second read aligns to the reverse DNA strand and the first read aligns to the forward DNA strand with the leftmost alignment position of the second read smaller than the leftmost alignment position of the first read. If the read pair is not extracted as discordant, Circle-Map will independently extract read pairs with any unaligned bases (soft-clipped and hard clipped).

### Realignment graph construction

Circular DNA identification begins by identifying clusters of candidate reads that are less than *K* nucleotides apart (discordant, soft-clipped and hard-clipped reads). For every cluster, we construct a weighted graph *G = (N,E)* with nodes *N = {n*_*0*_,*…,n*_*i*_*}*, which correspond to regions containing at least one breakpoint of unknown exact location, and edges *E = {e*_*0,*_*…, e*_*i*_*}*, which correspond to circle variants represented as connections between breakpoint regions (Fig. 1b). For every node n_i_, we obtain the set of edges connected to n_i_ by connecting the alignment positions of the reads belonging to node n_i_ with the alignment positions of their mates (for the discordant read pairs) and supplementary alignments (for the hard-clipped and soft-clipped reads). We then estimate edge weights 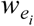 for every edge e_i_ using the mapping quality scores of the discordant mates and supplementary alignments that support the edge using:

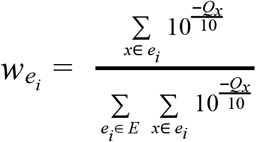

where *x* indicates a discordant read or supplementary alignment that supports e_i_ and *Q*_*x*_ indicates the phred scaled mapping quality of read *x*. Edges with weight below 0.01 were removed before the realignment step.

To define the final search space for the realignment step, node intervals are extended to [-*μ* - 5σ, *μ*+5σ], where *μ* and σ denote the mean and standard deviation of the insert size distribution estimated from concordantly mapped reads, to ensure that nodes originating from discordant reads will contain the breakpoint.

### Soft-clip realignment

In order to realign non-aligning bases of soft-clipped reads and obtain the circular DNA breakpoints at nucleotide resolution, we realign the soft-clipped parts of the reads probabilistically to the pruned realignment graph (Fig. 1c). We build a probabilistic model of the alignment using a Position-Specific Scoring Matrix (PSSM) that takes into account alignment mismatches and indels caused by sequencing errors.

Our algorithm begins by obtaining all possible alignments to the realignment graph using the infix Myers bit-vector algorithm as implemented in the Edlib library ^7,8^. In practice, we obtain the top scoring alignments to the graph by aligning the read and then masking the alignment coordinates, in order to keep searching for suboptimal alignments. For every alignment, we construct a PSSM using the following error model for matches and mismatches:

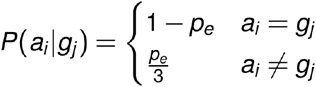

where *a*_*i*_ and *g*_*j*_ denote the identity of the bases on the read and the genome, respectively, and *p*_*e*_ indicates the probability of the base being sequenced wrong as determined from the base quality scores. Next, we compute the log-odds score for every base in the read by diving *P(a*_*i*_*|g*_*i*_) by the frequency of base *g*_*j*_ in the realignment graph, denoted as *q(g*_*j*_):

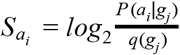

A score for the read is calculated by summing over the base scores in the PSSM:

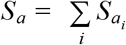

We then add an additional penalty to the PSSM score in order to account for insertions and deletions using an affine gap scoring scheme (adapted from ^9^) to yield the final alignment log-odds score:

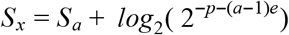

where *p* indicates the insertion and deletion penalty, *a* indicates the length of the event and *e* is the affine gap penalty.

Finally, Circle-Map converts the estimated alignment scores to alignment probabilities using:

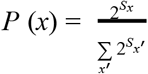

where the summation in the denominator runs over all possible alignments in the realignment graph. We consider a soft-clipped read realigned to the graph if the probability for its highest scoring alignment is greater than or equal to 0.99. The final realignment coordinates define putative circle coordinates (Fig. 1d). We consider a circular DNA as called if it is supported by a minimum of two breakpoint reads and contains at least one split read. Circle-Map reports all called circles with coordinates (chromosome, start and end), number of supporting reads (discordant and soft-clipped) and concordant coverage metrics (Supplementary methods).

We evaluated the performance of Circle-Map on both simulated data and real, circle-enriched data from human muscle tissue ^3,9^. As no publicly available tools for identification of circular DNA currently exist, we instead compared the performance of Circle-Map with that of CIRCexplorer2 ^10^, which is ranked as one of the best choices for circular RNA detection ^11^. Importantly, CIRCexplorer2 does not use splice-site or reference transcriptome information in the split read detection step. We further included Circle-Map without the realignment step as a baseline.

In the simulation benchmark, we simulated both high (30X) and low (7.5X) coverage sequencing of 13,097 circular DNAs across different lengths (range: 150-10,000 nts) and including SNVs and indels based on reference data from the 1000 genomes projects ^12^. To evaluate performance, we measured sensitivity, defined as the number of called circles present in the simulation set and precision defined as the fraction of correctly called circles found on the simulated set. On high coverage data (30X), Circle-Map attained a sensitivity of 0.945 by detecting 12,402 of the simulated circles and outperformed CIRCexplorer2 by 13 percentage points (Fig. 1a). A high sensitivity was also found on low coverage data (7.5X) were Circle-Map detected almost 70% of all circles in contrast to CIRCexplorer2 that detected 61% (Fig. 1b). Circle-Map and CIRCexplorer2 both attained precision higher than 0.98 on both high coverage and low coverage data sets (Fig. 1c and Fig. 1d).

We assessed the performance on real data using a paired-end sequencing dataset from a previous study ^3^, where circular DNA had been enriched from human muscle tissue by removal of linear DNA and amplification of the circular DNA prior to sequencing. As we lack a ground truth on real data, we instead used the fact that the data were enriched for circles and hence that most coverage from concordantly and contiguously aligned reads (i.e. non split-reads) across the genome should originate from circular DNA. We used the circle coverage fraction as an orthogonal proxy for correctness of the circle. Circle-Map detected 2,180 circles with >80% coverage while only 116 potential circles had a coverage less than 80% (Fig. 2e). In comparison, CIRCexplorer2 detected only 1,655 DNA circles with a coverage >80% and also detected a larger number of circles with coverage less than 20% (289 circles).

**Figure 2.**
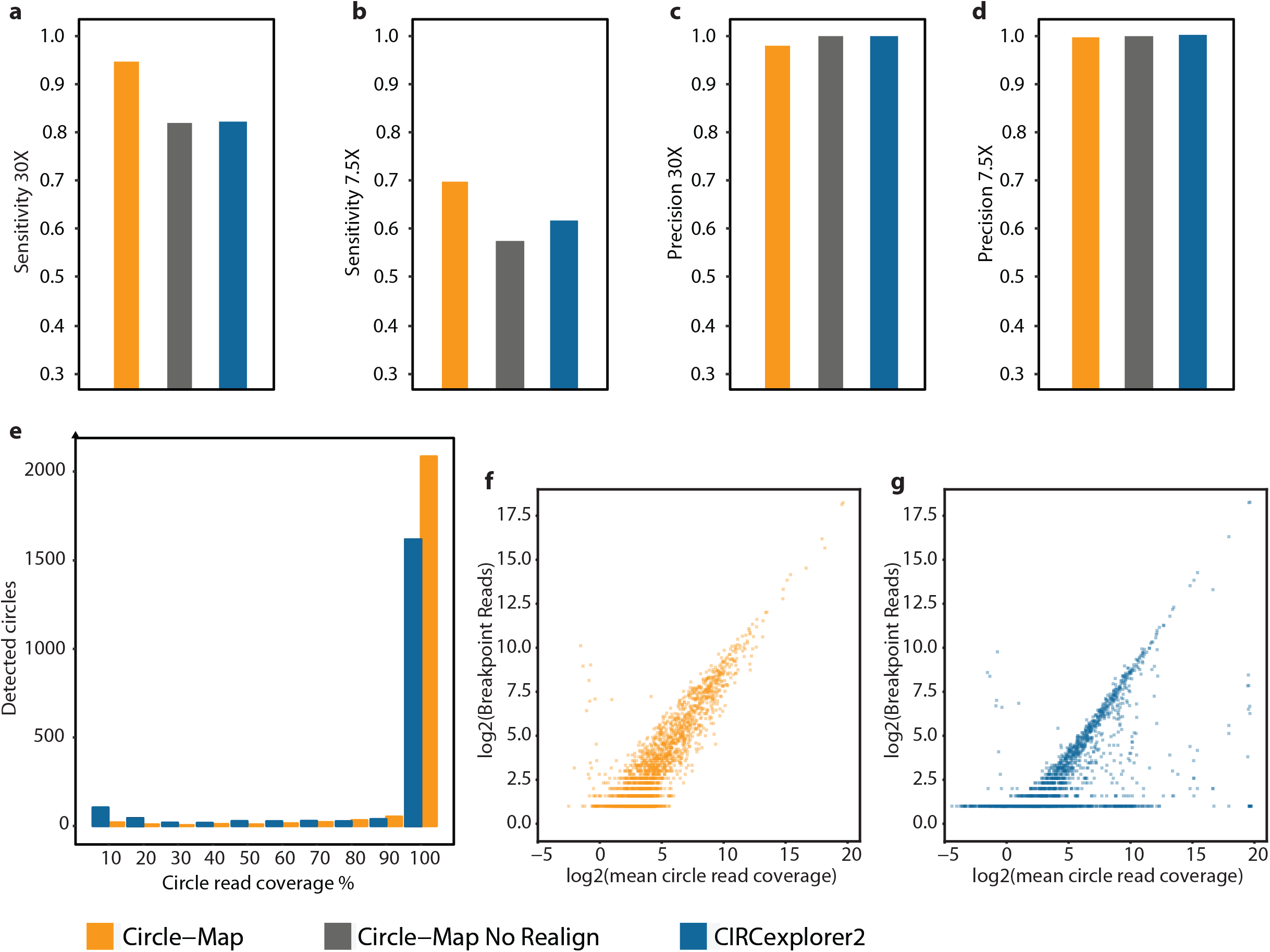
Evaluation of the circular DNA detection methods. **(a-d)** Circle-Map (orange), Circle-Map with no realignment (grey) and CIRCexplorer2 (blue) were evaluated on simulated circular DNA datasets with varying sequencing depths (e-g) and on real circle enriched data from human muscle. **(a)** Sensitivity at 30X and **(b)** 7.5X measured as the number of called circles found in the simulation set divided by the total number of simulated circles. Precision at **(c)** 30X **(d)** and 7.5X measured as the number of correctly called circles divided by the total number of called circles, true and false. **(e)** Histogram with the percentage of bases covered by sequencing reads for every circular DNA detected. The number of breakpoint reads (e.g. split and discordant reads) relative to the mean sequencing coverage within the circular DNA coordinates for **(f)** Circle-Map and **(g)** CIRCexplorer2.

Finally, we investigated the relationship between the number breakpoint reads (i.e split and discordant reads) and the concordant reads within the circle coordinates. We reasoned that in circle enriched data the breakpoint reads should correlate with the abundance of the concordant reads, so that strong disagreements between breakpoint and concordant reads should be indicative of erroneous circle detections (circles containing split reads with no coverage within the coordinates or circles containing few split reads in regions with high coverage). Hence, a plot of the number of breakpoint reads against the circle mean coverage can serve as a diagnosis tool to verify the agreement between breakpoint and concordant reads. Circle-Map obtained a Pearson’s correlation coefficient of 0.868 and showed a strong concordance between breakpoint reads and concordant reads (Fig. 2f). In contrast, CIRCexplorer2 achieved correlation of correlation of 0.582, with numerous data points substantially deviating from the central data trend Fig. 2g). Taken together, these data indicate that Circle-Map has both higher sensitivity and precision than CIRCexplorer2 in both simulated and real datasets.

In conclusion, we have developed a new method for detection of circular DNA based on a full probabilistic model for aligning reads across the breakpoint junction of the circular DNA structure. Using this model in combination with our guided realignment procedure, we are able to accurately align even very short soft clips (>5 nts). Using both simulated and real datasets, we have shown that this approach is both highly sensitive and precise - and significantly better than state-of-the-art methods from the related field of circular RNA detection, when applied to DNA. We predict that Circle-Map will find widespread usage across the expanding field of circular DNA research spanning from healthy and cancerous human tissues to model organisms such as yeast, worm and mouse. Finally, we believe that Circle-Map will likely also improve upon the state-of-the-art methods for detection of circular RNA.

## Supplementary methods

### Circle-Map simulation tool

In order to create a synthetic benchmark dataset we created a circular DNA simulation tool, included in the Circle-Map package. Our tool requires the genome reference sequence indexed with SAMtools ^13^ and the number of reads to simulate, and it will produce the simulated FASTQ files together with a BED file indicating the chromosomal coordinates of the circular DNA. The procedure involves three parts to simulate circular DNA. First, to consider genetic variation not present in the linear reference genome, it will introduce base substitutions and indels with BBMap (https://sourceforge.net/projects/bbmap/). Then, it will select the circle coordinates and the length from a uniform distribution (default range: 150-10000 nts) and sample the reads from the mutated genome. To generate all kinds of ordinary and circular DNA candidate reads our tool begins by defining the start and end read alignment coordinates and the insert size generated from a user defined normal distribution. Then, if the coordinates directly overlap the defined breakpoint coordinates, It will obtain both anchors of the read and join them together to generate the split read. Likewise, if the reads do not overlap the breakpoints but the insert within the paired reads spans the circle breakpoint it will generate a discordant read pair by sampling both reads from each side of the breakpoint. If any of the above mentioned conditions are not met, Circle-Map will sample a regular concordant read pair. Finally it will introduce instrument specific sequencing errors on the reads with ART ^14^.

### Simulated data

We used Circle-Map Simulate to generate 13097 circular DNA from the canonical chromosomes of the hg38 version of human genome, excluding the gap regions, downloaded from the UCSC genome browser ^15^ on the 1^st^ of February, 2019. We simulated the circular DNA with a circle length distribution ranging from 150 to 10000 nts. Altogether, we generated 2×10007326 paired end reads with a normally distributed insert size (mean = 300, s.d = 25) and a read length of 100 nts. We introduced indels and base substitutions on the reference at rates of 0.0001 and 0.001 respectively. Then, we generated reads with an Illumina HiSeq 2500 error profile without masking any bases (-nf option set to 0) and leaving the rest of the parameters as suggested by the ART authors.

We aligned the simulated reads to the canonical chromosomes of the canonical hg38 assembly using BWA-MEM^16^ (v0.7.17-r1188) without modifying the mapping qualities of the supplementary alignments (-q option turn on) and we used SAMtools^13^ (v1.9) to perform all the post processing step of the alignment files. In order to remove low quality alignments, we removed split reads were any of the split anchors had mapping qualities below 20 for both segments for CIRCexplorer2^10^ (v2.3.5), and we implemented the same filter internally on Circle-Map. Afterwards, we executed CIRCexplorer2 and Circle-Map leaving all the parameters as default and we removed the circular DNA containing less than two breakpoint reads, requiring Circle-Map to contain at least 1 split read in the set of the 2 breakpoint reads. Finally, as benchmark criteria for the simulated dataset, we used sensitivity, defined as the fraction of predicted circular DNA overlapping the simulated set by a fraction of 0.95, and precision, defined as the fraction of simulated circular DNA overlapping the predicted set by a fraction of 0.95.

### Real data

We downloaded a dataset (SRA ID: SRR6315430) from human muscle were the circular DNA was enriched prior to sequencing using the Circle-Seq^17^ procedure. Briefly the Circle-Seq procedure purifies DNA in 4 steps: isolation of the DNA by column separation, elimination of the residual linear DNA using exonuclease digestion followed by rolling circle amplification and paired sequencing on a Illumina HiSeq 2500 machine. In total, the dataset consisted of 2×12829402 paired reads with a length of 100 nts. We used prefetch and fasterq-dump from the SRA toolkit (https://github.com/ncbi/sra-tools v2.9.1) to download and convert the SRA files to FASTQ, respectively. We used the same strategy as described for the simulated data to detect the circular DNA in the real dataset. As performance measure for the real dataset, we first plotted a histogram representing the number of circular DNA against the percentage of bases covered within the detection coordinates of every circular DNA. We set the bins of the histogram to intervals of 10%. Finally, as second performance measure, we plotted the number of discordant reads and split reads against the mean sequencing coverage within the circular DNA coordinates, and we calculated the Pearson’s correlation coefficient as implemented in SciPy^18^.

### Circle-Map default circular DNA filtering

Circle-Map implements hard filters at the alignment and interval level to control for circle detection errors not accounted by the probabilistic model. At the alignment level, Circle-Map removes BWA-MEM flagged discordant reads pairs and split read primary alignments with mapping qualities below 20. The secondary alignments of the split reads with mapping qualities below 20 are not removed, but remapped using Circle-Map realignment algorithm in order to consider them as supportive for circular DNA. Furthermore, under Circle-Map realignment model, realigned reads with an edit distance greater than 0.05 as fraction of the read length are not considered. Circle-Map applies two hard filters to the detected intervals. First, Circle-Map removes circular DNA with allele frequencies smaller than or equal to 0.1, calculated as the number of split reads crossing the breakpoint divided by the mean sequencing coverage at the breakpoint nucleotides. Finally, to avoid redundant circular DNA identifications, circular DNA overlapping reciprocally by a fraction of 0.95 are combined into one interval.

### Circle-Map default output

Circle-Map will provides a tab separated file containing all the detected circular DNA. For every circular DNA will provide information containing the circular DNA mapping and information about the sequencing coverage in the circular DNA coordinates. Regarding the mapping based metrics, Circle-Map provides the mapping coordinates (chromosome, start and end), breakpoint read support (discordant and split reads) and a circular DNA mapping score, calculated by multiplying the length of the split read by its mapping probability, and summing over all the scores for the split reads supporting a circle. In relation to the sequencing coverage information, Circle-Map provides the mean sequencing coverage within the detection coordinates, standard deviation of the sequencing coverage, the fraction of circular DNA bases not covered by sequencing reads and, finally, the sequencing coverage increase upstream and downstream the detection coordinates. We calculated the increase in coverage as the ratio between the number of reads aligned 100 nts inside the breakpoint boundaries of the circular DNA and the same region extended 200 nts downstream (for the left part of the breakpoint) and upstream (for the right part of the breakpoint).

### Code availability

Circle-Map is implemented in python3 with the computational bottlenecks accelerated through multiprocessing and just-in-time compilation with Numba^19^. Circle-Map can be easily installed through the Python Package Index and the bioconda project^20^. The source code is released under the MIT license and it is freely available at https://github.com/iprada/Circle-Map.

